# Human PARPs modify RNA nucleobases *in vitro* and in cells

**DOI:** 10.64898/2026.03.31.715305

**Authors:** Michael Musheev, Jonas Siefert, Wiwik Bauten, Mareike Bütepage, Bernhard Lüscher, Christof Niehrs, Karla L. H. Feijs-Žaja, Roko Žaja

## Abstract

ADP-ribosylation is known as a protein modification, yet recent studies have expanded the range of ADP-ribosyltransferase (ART) substrates to include nucleic acids. tRNA 2′-phosphotransferase 1 (TRPT1) and several PARP family members can modify the 5′-phosphate of single-stranded RNA. Here, we show that PARP10 and PARP15 extend this activity beyond the 5′-phosphate terminus and generate N3-ADP-ribosyl uracil and N1-ADP-ribosyl guanine bases. The base-linked ADP-ribosylation is reversed selectively by the macrodomain-containing hydrolase TARG1. In TARG1 knockout cells, N1-ADP-ribosyl guanine can be detected. Together, these findings establish guanine and uracil ADP-ribose as two novel nucleotide modifications and reveal PARP15 and TARG1 as an enzyme pair which can dynamically regulate guanine ADP-ribosylation in living cells.

## INTRODUCTION

ADP-ribosylation is a covalent modification best known for proteins but increasingly recognised on nucleic acids^1-3^. ADP-ribosyltransferases (ARTs) catalyse this reaction using nicotinamide adenine dinucleotide (NAD^+^) as donor of ADP-ribose (ADPr). ARTs fall into two families, the diphtheria toxin-like ARTDs, and the cholera toxin-like ARTCs. The 17 human ARTD members, collectively called PARPs, install either single ADPr units (MARylation) or polymeric chains (PARylation)^4^.

In mammalian nucleic acids, internal base modification by ADP-ribosylation is already established for DNA as PARP1-dependent PARylation occurs on N1-adenosine and N3-cytidine^5,6^. Thus, base-linked modification of mammalian nucleic acid is chemically and enzymatically accessible. For mammalian RNA, however, the evidence centres on terminal phosphate modification. TRPT1 (KptA/Tpt1) ADP-ribosylates the 5′-phosphate of RNA *in vitro*, and PARP10, PARP11, and PARP15 modify RNA harboring 5′- or 3′-monophosphates^7,8^. ADP-ribosylated RNA has since been detected in cells^9^. 5′-ADPr-capped RNA is neither translated nor degraded, distinguishing it functionally from canonical caps, which promote translation, and NAD^+^ caps, which trigger degradation^10,11^. The cap is removed by ARH3 and macrodomain hydrolases including MACROD2, PARG, and TARG1^12^.

Internal RNA ADP-ribosylation, by contrast, has so far been observed only in non-mammalian systems. The *Streptomyces coelicolor* transferase ScARP modifies guanosine at the N2 position within tRNA ^13^, and pierisin toxins ADP-ribosylate the same position in double-stranded DNA^14^. Guanine-base ADPr is reversed by the bacterial and archaeal NADAR family^15^. A chemically distinct modification is installed by the Pseudomonas aeruginosa toxin RhsP2, which ADP-ribosylates the 2′-OH of double-stranded RNA, inhibiting translation^16^. 2′-OH ADPr is reversed by bacterial and human PARG but not by other macrodomain hydrolases or NADAR enzymes^15^, highlighting that different linkage chemistries require different erasers.

Critically, no mammalian PARP has been shown to ADP-ribosylate internal RNA nucleobases, no such marks have been detected in mammalian cells, and it is unknown which hydrolases would erase them. Here we address these gaps by investigating PARP15, for which we previously showed that overexpression in HeLa cells increases levels of ADPr-RNA^9^. We now report that in addition to modifying the 5’-phosphate, PARP10 and PARP15 also modifies guanosine and uracil bases. This modification cannot be reversed by the majority of ADP-ribosylhydrolases but is efficiently removed by TARG1. The relevance of TARG1 for this modification is underlined by our finding that ADPr-guanosine is detectable in diverse human cells upon TARG1 knockout only. Our work establishes N3-ADPr-U and N1-ADPr-G as novel RNA modification.

## RESULTS

### PARP15 modifies RNA beyond the 5′ phosphate through a noncanonical ADP-ribose linkage

Overexpression of PARP10-12 or PARP15 increases cellular ADPr-RNA levels, with PARP15 showing the strongest induction^9^. The identity of PARP15 substrates remains unknown, and therefore we analysed its catalytic activity. Using a fixed concentration of a 5′p-RNA-3’Cy3 oligonucleotide and increasing enzyme amounts, we found that TRPT1 produces a single extra band, whereas catalytic domain of PARP15 (PARP15cat) generated multiple bands, consistent with the addition of several ADPr units (**Figure 1A**). Increasing reaction time induced these extra bands further (**Figure S1A**), whereas the catalytic mutant H559Y was not active and modification was reduced by the dual PARP10/15 inhibitor OT-42 (**Figure 1B**)^17^. This confirms the activity is intrinsic to PARP15. This incorporation of multiple ADPr units leads to two possible scenarios: (1) PARP15 generates poly(ADPr) at the 5′ phosphate, or (2) it installs multiple ADPr moieties at internal RNA positions (**Figure 1C**). PARG is a poly(ADP-ribosyl)glycohydrolase^18^ and is capable of removing phosphoester-linked ADPr from RNA^7^, and thus useful to test if scenario 1 holds true. Incubation with PARG released only a single ADPr from PARP15-modified RNA, while demodifying TRPT1-modified substrates (**Figure 1D**), indicating that PARP15 adds multiple independent ADPr adducts rather than poly(ADP-ribose) chains. The de-AMPylase aprataxin (APTX), also shifted PARP15-modified RNA, albeit less efficiently than PARG, suggesting selective removal of the AMP from several ADPr-RNA linkages rather than PAR. To confirm the presence of additional PARP15 acceptor sites on RNA we tested activity on 5′p and 5′OH oligonucleotides. PARP15cat modified both, whereas TRPT1 requires a 5′ phosphate (**Figure 1E**). PARP15 activity on 5′OH RNA increased with enzyme concentration and time and was lost with the H559Y mutant or in the presence of PARP inhibitor (**Figure S1B and Figure S1C**). Additionally, we acid-treated the ADPr-RNA oligos, as 5′ phosphoester bonds are acid-labile. Indeed, TRPT1-modified RNA at the 5’-p lost the ADPr moiety upon acid treatment (**Figure S1D**), while PARP15-modified 5’OH RNA was not acid sensitive (**Figure S1E**).

**Figure 1.**
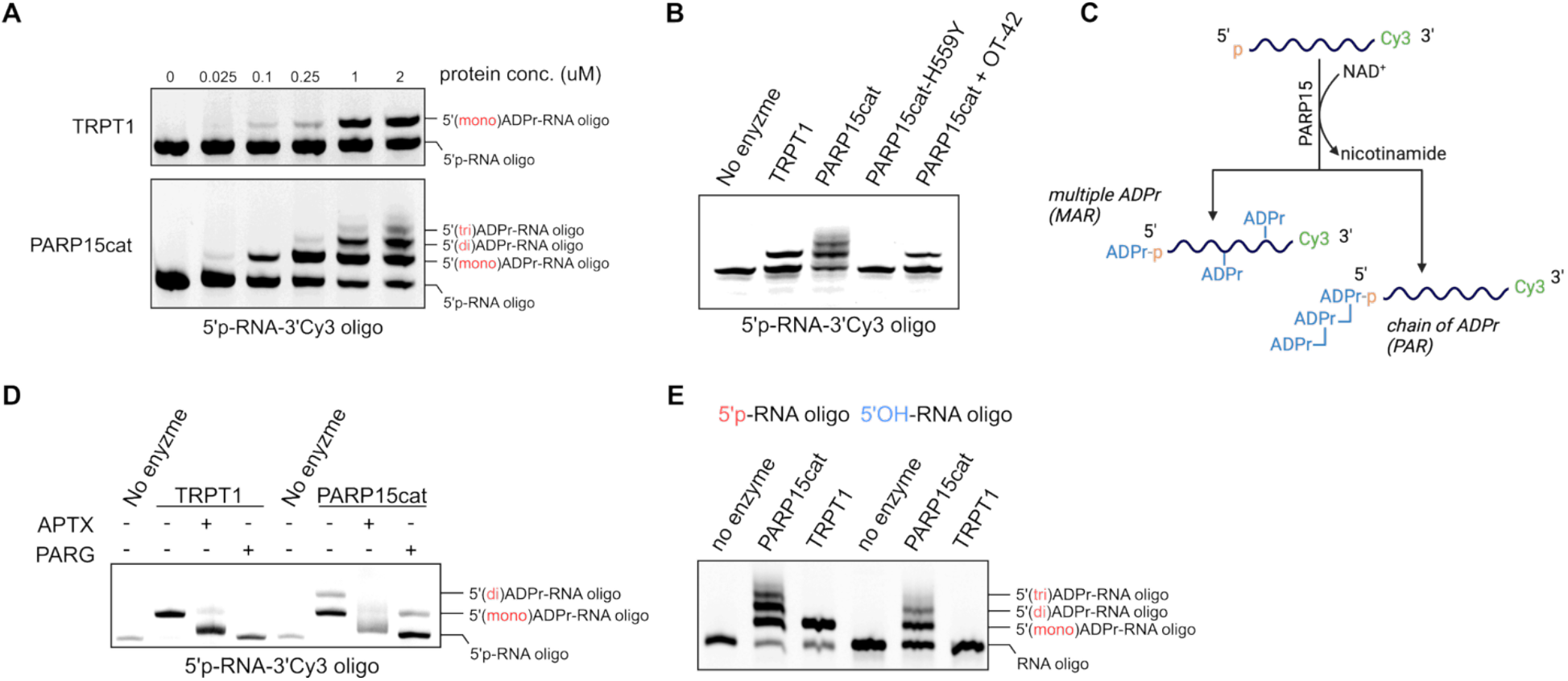
PARP15 modifies RNA via a noncanonical ADP-ribose linkage. **(A)** PARP15cat and TRPT1 were incubated with a 5’-p RNA-Cy3 in presence of 500 µM NAD^+^. The oligonucleotides were separated using urea-PAGE and visualised using in-gel fluorescence. **(B)** *In vitro* ADP-ribosylation assay as in (A), with a PARP15-H559Y mutation or with addition of 1 µM PARP15 inhibitor OT-42. **(C)** Schematic overview of the possible RNA oligonucleotide modification by PARP15. **(D)** A 5’-p ssRNA-Cy3 oligo was incubated with TRPT1 or PARP15cat in presence of 500 µM NAD^+^, purified and incubated with APTX or PARGcat. Reactions were analysed using urea-PAGE and in-gel fluorescence. **(E)** PARP15cat and TRPT1 were incubated with a 5’-p ssRNA-Cy3 or 5’-OH ssRNA-Cy3 oligonucleotide in presence of 500 µM NAD^+^. Reaction products were analysed using urea-PAGE and in-gel fluorescence.

Together, these data indicate that PARP15 installs multiple ADPr units on RNA via two distinct linkages: an acid-labile phosphoester at the 5′ monophosphate and a PARG-resistant, acid-stable bond internally.

### PARP15cat modifies guanine and uracil bases of RNA

In other organisms, ADP-ribosylation of nucleobases occurs, where for example ScARP ADP-ribosylates the guanosine base in RNA and DNA^13^ and the ribose 2’-hydroxyl group of RNA is ADP-ribosylated by RhsP2^16^. We thus hypothesized that PARP15 can ADP-ribosylate RNA on either the base or 2’-hydroxyl group of ribose on RNA. To test whether 2’-hydroxyl group of ribonucleotides serves as acceptor site, we treated the PARP15cat modified oligo with RNase 4. RNase 4 is a single-strand endoribonuclease that cleaves RNA at uridine-purine dinucleotide sites, however, activity is inhibited by ribose modificatins such as 2’-O methyluridine, whereas uridine base modifications have no effect on RNase 4 activity^19^. We observed that both a PAP15cat modified oligo and non-modified control RNA oligo were digested by RNase 4 (**Figure 2A**). This indicates that PARP15 most likely modifies the base.

**Figure 2.**
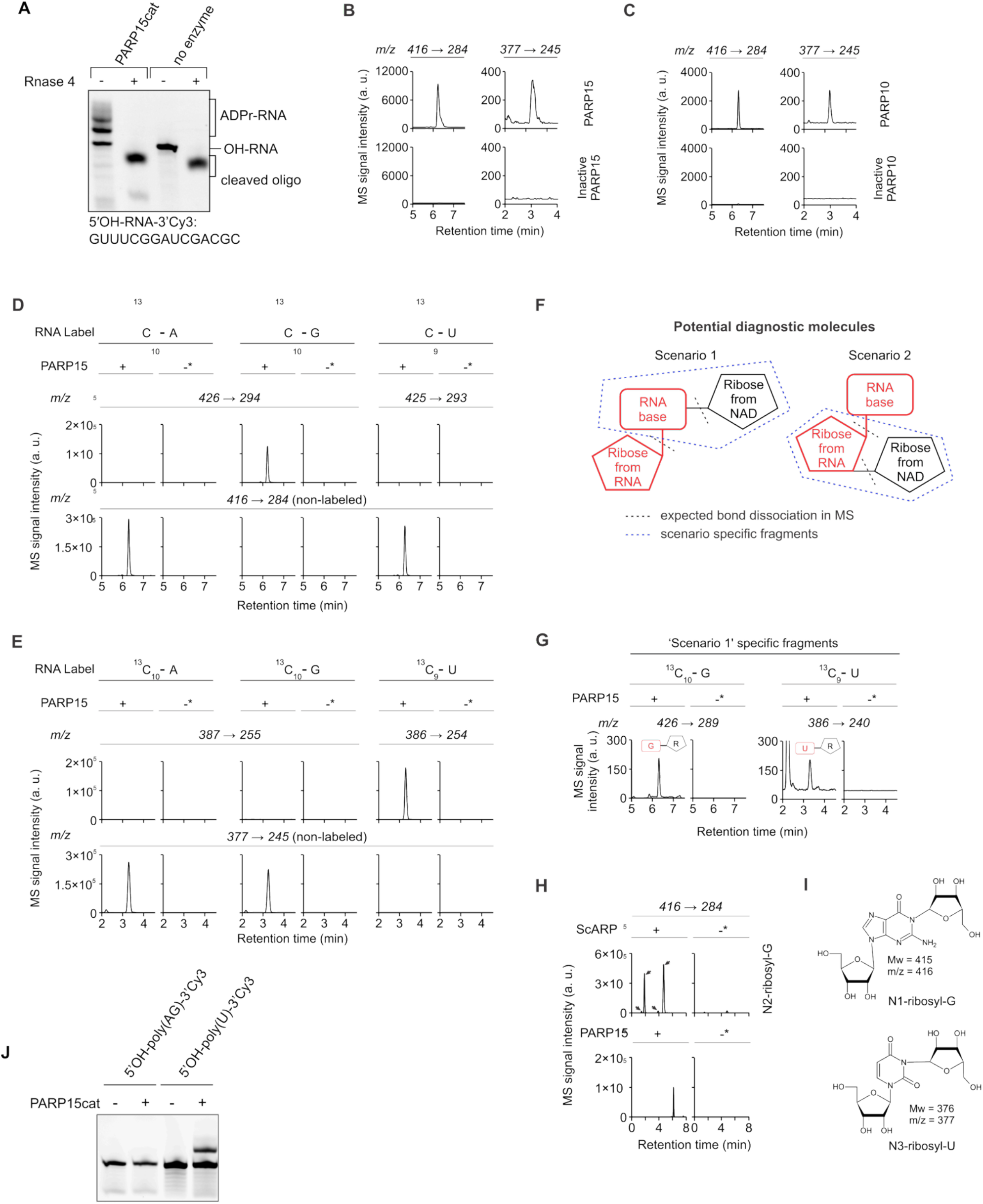
PARP15cat modifies the guanine and uracil base. (**A**) A PARP15cat ADP-ribosylated 5’-OH ssRNA-Cy3 or non-modified control were incubated with RNase 4 for 20 min at 37ºC. Reactions were separated using urea-PAGE and in-gel fluorescence was detected. (**B**,**C)** LC-MS/MS chromatograms of reaction products from *in vitro* ADP-ribosylation assays in presence of native or heat-denatured PARP15cat (B) or PARP10cat (C). **(D**,**E)** LC-MS/MS chromatograms of reaction products from *in vitro* ADP-ribosylation assays in presence of native or heat-denatured PARP15 with three isotopically labeled RNA substrates as indicated. Top, scan for signals with m/z transitions expected for a mass shift of ‘nucleoside 416’(C) and ‘nucleoside 377’(D) from each label. Bottom, detection of signals that correspond to non-labeled ‘nucleoside 416’ (C) and nucleoside 377’ (D). **(F)** A diagram indicating two diagnostic molecules generated after enzymatic degradation of ADPr-RNA from two scenarios: 1-ADP-ribosylated at the base; 2-ADP-ribosylated at the RNA ribose, with bonds most susceptible to fragmentation indicated by dashed lines, and the encircled parts of the molecules indicating scenario-specific fragments. **(G)** LC-MS/MS chromatograms of products from (C) and (D), detecting scenario 1-specific fragment from (E). **(H)** LC-MS/MS chromatograms of reaction products from *in vitro* ADP-ribosylation assays with either ScARP or PARP15cat. Arrows indicate reaction-specific products. **(I)** Structures of diagnostic molecules deduced from LC-MS/MS experiments, indicating the N1 position of G and N3 position of U as PARP15 targeting sites for attachment of ADP-ribose. **(J)** PARP15cat was incubated with 5′OH-poly(U)-3′-Cy3 (U-only) or 5′OH-poly(AG)-3′-Cy3 (mixed G/A) oligonucleotides in the presence of 500 µM NAD^+^; reactions were analysed by urea-PAGE and in-gel fluorescence.

To identify the site of modification, a PARP15cat modified RNA oligonucleotide was enzymatically digested to its constituent nucleosides. This releases unmodified nucleosides or when ADP-ribosylated, release a ribose-linked nucleoside. The resulting products were analysed by LC-MS/MS to monitor for novel peaks corresponding to the mass addition of a ribose to any nucleoside atom. We observed two new peaks with a positively charged ion m/z of 416 (‘nucleoside 416’) and 377 (‘nucleoside 377’), matching to an addition of ribose to either guanine or uracil nucleotides respectively. These peaks were absent when heat-inactivated PARP15cat was used (**Figure 2B**). When testing additional PARPs, we noticed that also PARP10 activity leads to similar patterns, indicating that this activity is not unique to PARP15 (**Figure 2C**). To confirm that the new signals arose from ADP-ribosylated guanine or uracil, we used three RNA substrates, each containing a single isotopically labelled canonical nucleotide (^13^C_10_-G, ^13^C_10_-U or ^13^C_10_-A). Only the ^13^C_10_-G containing substrate shifted the m/z 416 signal to 426 (**Figure 2D**), as expected for guanosine labeling. Analogous patterns were observed for ‘nucleoside 377’, except that it showed a labelled uridine-specific signature (**Figure 2E**). These experiments prove that ‘nucleoside 416’ is ribosyl-guanosine; and nucleoside 377’ is ribosyl-uridine. To determine whether the NAD^+^-derived ribose is attached to the base or ribose moiety, we exploited the distinct MS/MS fragments predicted for each structure (**Figure 2F**, dashed lines). Isotopically labelled nucleosides allowed us to distinguish the heavy ribose contributed by the nucleotide from the natural ribose derived from NAD^+^. Detection of fragments unique to base attachment therefore directly identified base-linked ADP-ribosylation. Such fragments were observed for both U- and G-ADP-ribosylation products, indicating that the modification occurs on the base (**Figure 2G**), corroborating the RNase 4 digest. This finding indicates that ADPr is attached to uridine at N3 and to guanosine at either N1 or N2. ScARP was reported to modify guanosine at N2 position previously ^13,20^. We performed ADP-ribosylation reactions with ScARP and PARP15cat and compared the products in LC/MS. ScARP-mediated N2-ribosyl-G elutes earlier (**Figure 2H**) and has a signature with several peaks of identical mass eluting at different times. This result leaves one possible structure: N1-ribosyl-G. Thus, PARP15 ADP-ribosylates both G and U on their N1 and N3 positions respectively, which after degradation, release diagnostic molecules N3-ribosyl-U and N1-ribosyl-G (**Figure 2I**). To confirm the LC/MS analysis we incubated different oligonucleotides with PARP15cat and observed efficient ADP-ribosylation of an oligonucleotide containing exclusively uracil, confirming that PARP15cat can modify this nucleotide (**Figure 2J**). However, guanosine in a GA-containing RNA oligo was not modified, indicating that guanosine may need a specific motif for modification.

### Molecular basis for TARG1 hydrolysis of guanine-N1-ADPr and uracil-N3-ADPr

We next asked which enzymes remove base-linked ADPr. We purified 5’-ADPr-poly(U)-3’Cy3 and tested a panel of hydrolases (**Figure 3A**). ARH3, MACROD2, and PARG did not remove the modification even at a concentration of 1 μM enzyme whereas TARG1 efficiently hydrolysed uracil-N3-ADPr. The same pattern of hydrolysis was observed on 5′ADPr-OH-RNA-3’Cy3 (**Figure S2**) where the completeness of the reversal indicated that TARG1 hydrolyses uracil-N3-ADPr and guanine-N1-ADPr. To confirm TARG1 specificity toward the N1 but not N2 guanine linked ADPr, we modified an RNA oligo using ScARP (N2-ADPr) or PARP15cat (N1-ADPr) and incubated these with TARG1 or NADAR. NADAR exclusively hydrolysed guanine-N2-ADPr but not guanine-N1-ADPr (**Figure 3B**). By contrast, TARG1 was not active on guanine-N2 linked ADPr but fully hydrolysed the PARP15cat modified oligo demonstrating its specificity toward guanine-N1-ADPr (**Figure 3C**). We confirmed this observation by analysing the same reactions using LC-MS/MS (**Figure S3A-B**). These results demonstrate TARG1 specificity for the novel RNA modification guanine-N1-ADPr.

**Figure 3.**
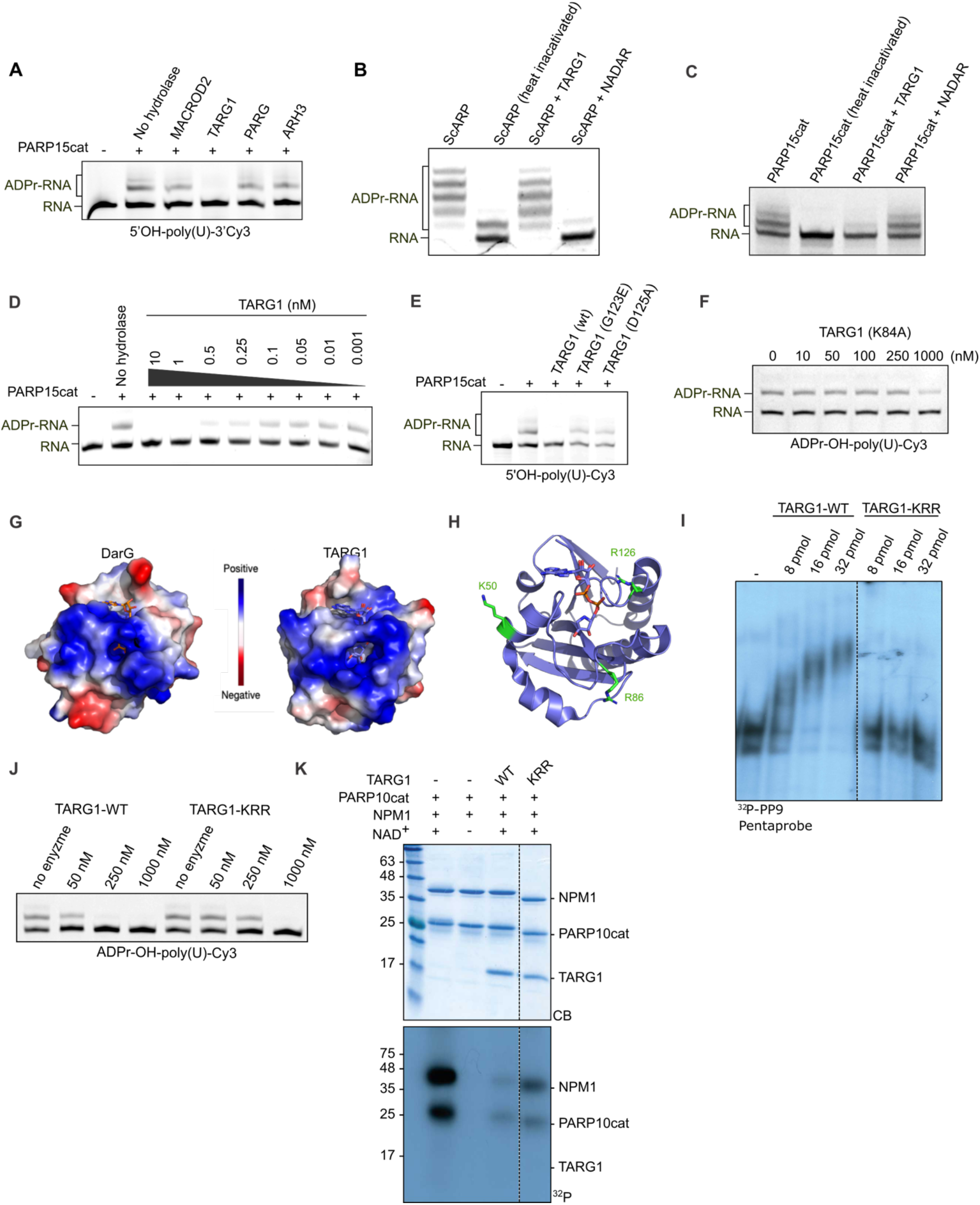
Molecular basis for TARG1 hydrolysis of guanine-N1-ADPr and uracil-N3-ADPr. **(A)** A poly(U)-Cy3 oligo was ADP-ribosylated by PARP15cat and purified after proteinase K digest. Activity of different hydrolases was assessed by incubating the ADPr-RNA oligo with 1 µM of indicated hydrolases for 30 min at 37ºC. Reactions were resolved on urea-PAGE and in-gel fluorescence detected. **(B)** Oligonucleotides were ADP-ribosylated with ScARP and purified as in (A) and incubated with either 100 nM NADAR or TARG1 for 30 min at RT. After separation on urea-PAGE the gels were stained with SYBR Gold and analysed by in-gel fluorescence. **(C)** Oligonucleotides were ADP-ribosylated with PARP15cat and ADPr-ribosylhydrolase activity of 100 nM NADAR and TARG1 analysed as in (A). **(D)** ADPr-RNA-poly(U)-Cy3 was prepared using PARP15cat as in (B) and then incubated with indicated concentrations of TARG1, followed by urea-PAGE and in-gel fluorescence detection. **(E)** As in (D), with inclusion of TARG1 mutants at 50 nM concentration. **(F)** ADPr-poly(U)-Cy3 prepared as in (D) was incubated with indicated concentrations of TARG1 K84A mutant and detected as in (D). **(G)** Comparison of the protein surfaces of DarG (5m3e) and TARG1 (4j5r), highlighting the electrostatic surface potential. **(H)** Cartoon representation of TARG1 crystal structure (4j5r) with indicated residues contributing to the positively charged surface potentially involved in RNA binding. **(I)** Electrophoretic mobility shift assay with ^32^P-labelled pentaprobe RNAs. Purified pentaprobe 9 (^32^P-PP9) was incubated with the indicated amounts of TARG1 or TARG1 triple K50D/R86D/R126D mutant (KRR). Unbound RNA was separated from protein-bound RNA on native 7% polyacrylamide gels. Mobility shifts were analysed by autoradiography. **(J)** ADPr-poly(U)-Cy3 was prepared as in (A) prior to incubation with indicated amounts of TARG1 wild-type or K50D/R86D/R126D mutant (KRR). Reactions were separated using urea-PAGE and in-gel fluorescence was detected. **(K)** A model substrate protein, NPM1, was ADP-ribosylated using PARP10cat and radiolabelled NAD^+^. ADP-ribosylated proteins were incubated with TARG1 or the indicated mutants. Reactions were separated using SDS-PAGE, followed by Coomassie staining (CB) and detection of ^32^P by radiography.

TARG1 hydrolyses uracil-N3-ADPr in the low-nanomolar range (**Figure 3D**), making it at least an order of magnitude more efficient than on ADP-ribosylated protein substrates or on 5′-ADPr-pRNA^12^. Although TARG1 adopts the conserved macrodomain fold, it uses a distinct catalytic mechanism. TARG1 operates via a lysyl-ADPr covalent intermediate, reminiscent of OGG1-family DNA glycosylases^21^. Lys84 acts as the nucleophile and attacks the C1″ position of ADP-ribose, generating a transient Lys84-ADPr adduct. This intermediate is subsequently hydrolysed by a water molecule activated by Asp125 to release ADPr. Consistent with this model, both TARG1 D125A and K84A mutants exhibited impaired hydrolysis of U-N3-ADPr (**Figures 3E and 3F**). This two-step “glycosylase-like” mechanism may be well suited for cleaving the N-glycosidic uracil-N3-ADPr linkage, because TARG1 first captures ADPr covalently before resolving the Lys-ADPr bond, rather than relying on a single direct water-attack step.

Similarly to DarG, which hydrolyses guanine-N2-ADPr^13,20^, TARG1 displays a positively charged surface around the catalytic centre (**Figure 3G**). To test whether this surface contributes to ADPr-RNA docking, we purified the triple mutant TARG1 K50E/R86D/R126D (“KRR”) (**Figure 3H**). The KRR mutant TARG1 displayed reduced RNA binding (**Figure 3I**) as well as reduced hydrolase activity on uracil-N3-ADPr (**Figure 3J**). In contrast, the protein was active on ADPr-protein (**Figure 3K**), indicating that the reduced activity is not due to protein misfolding.

### Guanine is ADP-ribosylated at N1 in human cells

PARP10 and PARP15 can thus modify nucleobases *in vitro*, leading to modification at heterocyclic nitrogen N3 or N1 in uracil and guanine, respectively (**Figure 4A**). The next question is whether this modification also occurs in vivo. We observed that PARG inhibition is needed to detect ADPr-pRNA^12^ and therefore generated a TARG1 CRISPR/Cas9-mediated knock-out in HeLa cells to enhance LC-MS/MS detection of ADPr-bases. We did not detect guanine-N1-ADPr in wild-type cells, however upon PARP15 overexpression in TARG1-KO cells a peak corresponding to guanine-N1-ADPr was observed (**Figure 4B**). Our *in vitro* PARP15 data showed that U and G bases can be modified but we did not observe any U-N3-ADPr in cellular RNA (**Figure 4C**). To obtain a more quantitative measure of the amount of base-modified RNA we incubated isolated cellular RNA with TARG1 to release ADPr, followed by incubation with NUDT5. NUDT5 processes the free ADPr to form AMP and ribose-phosphate, which can be used to determine AMP levels as proxy for ADPr-RNA levels. We used RNA oligos modified with TRPT1 (ADPr-pRNA), ScARP (guanine-N2-ADPr) or PARP15cat (guanine-N1-ADPr or uracil-N3-ADPr) to test the method (**Figure 4D**). As expected, PARG released ADPr from ADPr-pRNA but not from either of the base modified oligos, NADAR specifically removed ADPr from guanine-N2-ADPr-RNA and TARG1 only reversed ADPr from guanine-N2-ADPr and uracil-N1-ADPr. We next used this assay to measure guanine-N1-ADPr in cellular RNA, extracted from wild-type or TARG1 knockout cells with or without overexpression of PARP15. A modest increase of released ADPr was observed in PARP15 overexpressing cells, whereas TARG1 knockout led to a strong increase in ADPr, confirming the presence of N1-ADP-ribosylated guanine in total cellular RNA (**Figure 4E**). To corroborate these findings, we isolated RNA from mammalian cell lines with TARG1 knockout or knockdown. Basal levels of the modified bases were low in wild-type cells, but increased upon TARG1 knockdown in all cell lines tested (**Figure 4F and 4G**).

**Figure 4.**
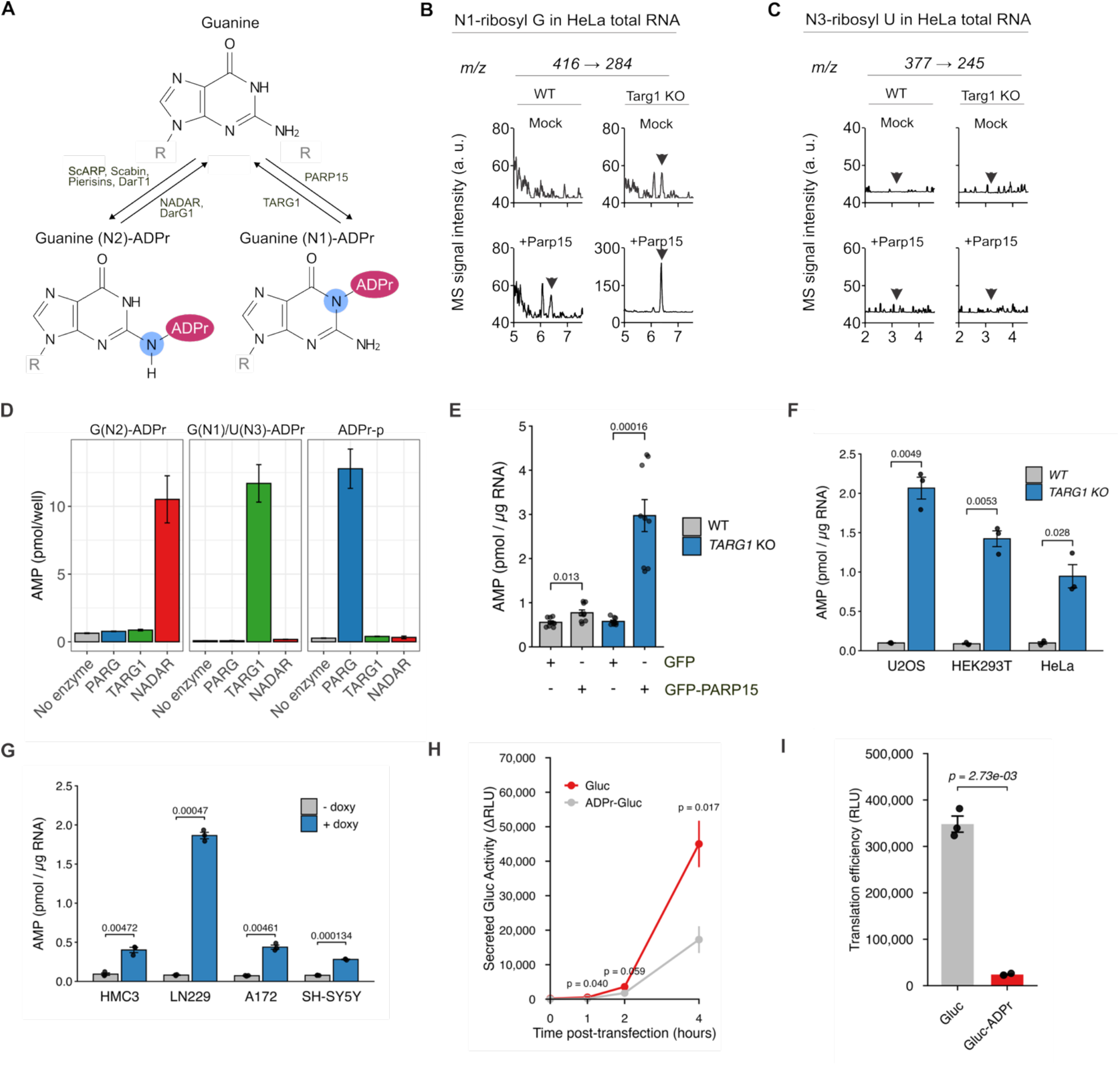
Guanine is ADP-ribosylated at N1 in human cells. **(A)** Overview of the guanine base modification as introduced by ScARP-type toxins or by mammalian PARPs. **(B)** LC-MS/MS chromatograms monitoring N1-ribosyl-G in digested total RNA from wild type (WT) and TARG1 KO HeLa cells overexpressing either EGFP alone (MOCK) or PARP15-EGFP (PARP15) fusion proteins. Arrows indicate the appearance of the peak corresponding to N1-ribosyl-G. **(C)** LC-MS/MS chromatograms monitoring N3-Ribosyl-U in digested total RNA from wild type (WT) and TARG1 KO HeLa cells overexpressing either EGFP alone (MOCK) or PARP15-EGFP (PARP15) fusion proteins. Arrows indicate the expected elution time for N3-ribosyl-U. **(D)** 5’OH-RNA oligos were ADP-ribosylated using ScARP or PARP15cat; a 5’-pRNA oligo was ADP-ribosylated using TRPT1. After purification ADP-oligos were incubated with indicated hydrolases (100 nM) at RT for 30 min. ADPr was released by incubation with 200 nM NUDT5 and AMP was measured using AMP-Glo. **(E)** RNA was isolated from HeLa TARG1 knock-out cells transfected with GFP or GFP-PARP15. Extracted RNA was first incubated with 100 nM TARG1 followed by NUDT5 and AMPGlo assay as in (D). **(F)** ADPr release from total cellular RNA isolated from indicated TARG1 knockout cell lines measured as in (D). **(G)** ADPr release measured as in (F) with total cellular RNA isolated from different TARG1 knockdown cell lines. TARG1 knockdown was induced in cells stably transfected with Tet-pLKO-puro plasmids containing TARG1 specific shRNA using 200 ng/ml doxycycline (72h). **(H)**. HEK293T cells were transfected with Gaussia luciferase (Gluc) mRNA modified by PARP15cat or control mRNA and medium was collected at indicated intervals. Translation of control Gluc mRNA (grey) is compared to ADPr-Gluc mRNA (red). Data represent baseline-subtracted translation efficiency (ΔRLU). Points represent the mean of n=4; error bars standard deviation. P-values indicate significance as determined by Welch’s t-test. **(I)** Endpoint analysis of *in vitro* translation efficiency measured after 4h incubation with non-modified Gluc mRNA and enriched ADPr-Gluc mRNA. Bars represent the mean translation efficiency (RLU) of n=3; and error bars represent standard deviation. Individual data points are shown as jittered dots. P-value indicates significance as determined by Welch’s t-test.

The bacterial anti-phage defense toxin CmdT functions as an mRNA-specific ADP-ribosyltransferase, attaching an ADPr moiety to the N6 position of adenines to arrest translation^22^. Therefore we tested whether the guanine-N1 ADP-ribosylation inhibits translation in human cells. We transfected an m^7^G-capped, poly(A) tailed Gaussia luciferase reporter mRNA modified with ADPr (ADPr-Gluc) and quantified secreted luciferase (**Figure 4H**). ADPr-RNA was translated to a lower extent, indicating that ADP-ribosylation of mRNA could block translation. We also measured translation efficiency using an *in vitro* translation system. Here we used a Gluc reporter with a ribosomal-binding site in the 5′ UTR and a stable hairpin in the 3′ UTR as a translation termination signal. Following *in vitro* transcription and modification by PARP15, we enriched the ADPr-RNA and measured translation. We observed a striking reduction in translation of the ADPr-Gluc mRNA (**Figure 4I**), which confirms that base-targeted ADP-ribosylation directly interferes with translation.

## DISCUSSION

Here we show that mammalian ARTs can ADP-ribosylate RNA nucleobases and that these modifications are reversed by TARG1, extending mammalian RNA ADP-ribosylation beyond terminal phosphates. These findings underscore that RNA is a bona fide ADP-ribosylation substrate in both prokaryotes and eukaryotes. *In vitro* data show that PARP10 and PARP15 modify RNA via two distinct linkages: an acid-labile 5′-phosphoester that can be processed by PARG, and an acid-stable, PARG-resistant linkage consistent with base modification. Using LC-MS/MS, we were able to precisely define the modification as guanine-N1-ADPr and uracil-N3-ADPr. Guanine-N1-ADPr is present in cellular RNA, establishing RNA base ADP-ribosylation as a novel epitranscriptomic mark in mammalian systems. The presence of RNA ADP-ribosylhydrolase TARG1, that selectively removes uracil- and guanine-linked ADPr, supports the view that RNA base ADP-ribosylation is a regulated modification. TARG1 deficiency causes a neurodegenerative phenotype via an unknown mechanism assumed to be connected to protein ADP-ribosylation^21^. An alternative possibility is that accumulation of ADPr-RNA, rather than ADP-ribosylated proteins, underlies this phenotype. In this model, failure to remove aberrant ADPr-RNA would compromise RNA metabolism and associated stress responses, providing an avenue to be investigated in regards to the TARG1^-/-^ phenotype. The essential nature of the ADPr-base eraser TARG1 does underscore the functional relevance of this novel RNA modification.

The discovery of nucleobase ADP-ribosylation in mammalian cells perhaps causes more questions than it answers. We have removed TARG1 to stabilise and detect the modification in cells, which raises the question which (patho)physiological conditions do the same? Is the RNA modification upregulated in response to a specific stress, similar to protein ADP-ribosylation, where DNA damage^23^, proteotoxic stress^24^ as well as viral infection^25^ impact on ADP-ribosylation? Assuming this to be the case, how does nucleobase ADP-ribosylation contribute to the cellular stress responses? Is its main function the recruitment of specific ADPr-nucleobase binding proteins to determine RNA fate? Or does it directly impact RNA function, and influence RNA metabolic processes such as degradation, splicing or translation? To answer these questions, two aspects will be of utmost importance for future studies: the identification of the readers of this novel modification, as well as the identification of the RNAs that are modified under specific conditions. Only after mapping of the proteins interacting with ADPr-RNA as well as characterising the nature of RNAs modified, can we start drawing models of the functional relevance of RNA nucleobase ADP-ribosylation.

In sum, our data establish that mammalian PARPs can generate G-N1-ADPr or U-N3-ADPr *in vitro* with detection of G-N1-ADPr in human cells, thereby adding a new class of reversible base-linked modification to the epitranscriptome.

### Limitations of the study

Several questions remain open: although we identified G-N1-ADPr upon PARP15 overexpression, other enzymes may install this modification in cells, as exemplified by *in vitro* modification of both guanine and uracil by PARP10. Second, the RNA sequence and structural motifs that license PARP15-dependent guanine modification are not yet defined. Third, the distribution and dynamic regulation of RNA base ADP-ribosylation remain unknown.

## RESOURCE AVAILABILITY

### Lead contact

Requests for further information and resources should be directed to and will be fulfilled by the lead contact, Roko Žaja.

### Materials availability

Plasmids generated in this study have been deposited to Addgene, with a list of plasmid names and Addgene catalog numbers available in the methods section. Stable cell lines are available from the lead contact with a completed materials transfer agreement.

### Data and code availability

This paper does not report any original code. Any additional information required to reanalyze the data reported in this paper is available from the lead contact upon request.

## ACKNOWLEDGMENTS

We would like to thank Michael Cohen and Lari Lehtiö for generously providing materials, Jinyu Li and Herwig Schüler for fruitful discussions about TARG1 structure, and Nils Hohaus for technical support. This work was funded by the Deutsche Forschungsgemeinschaft DFG [ZA 1248/3-1 to R. Ž.] and a habilitation stipendium from the Medical Faculty of RWTH Aachen University to K.F-Ž.

## AUTHOR CONTRIBUTIONS

Conceptualization, R.Ž.; methodology, M.M., K.F.Ž. and R.Ž.; Investigation, M.M., J.S, W.B., M.B., K.F.Ž and R. Ž.; writing-original draft, M.M., K.F.Ž. and R.Ž.; funding acquisition, K.F.Ž. and R.Ž.; resources, B.L. and C.N.; supervision, K.F.Ž. and R.Ž.

## DECLARATION OF GENERATIVE AI AND AI-ASSISTED TECHNOLOGIES

No generative AI or AI-assisted tools were used.

## SUPPLEMENTAL INFORMATION

**Document S1. Figures S1–S4, Tables S1-S3**

## STAR★METHODS

### KEY RESOURCES TABLE

| REAGENT OR RESOURCE | SOURCE | IDENTIFIER |
| --- | --- | --- |
| **Antibodies** |  |  |
| Rabbit polyclonal anti-TARG1 | ref | N/A |
| Mouse polyclonal anti-GFP | Rockland | 600-101-215, RRID: AB_218182 |
| HRP-Goat anti-Rabbit IgG (H+L) | Jackson ImmunoResearch | 111-035-144; RRID: AB_2307391 |
| Poly/Mono-ADP Ribose (D9P7Z) Rabbit Monoclonal Antibody | Cell signaling technology | 89190, RRID: AB_3716623 |
| **Bacterial and virus strains** |  |  |
| DH5α | New England Biolabs | C2987 |
| BL21(DE3) | New England Biolabs | C2527 |
| Rosetta<sup>TM</sup> | Novagen | 70954 |
| **Chemicals, peptides, and recombinant proteins** |  |  |
| TRIzol Reagent | Thermo Fisher Scientific | 15596026 |
| β-NAD<sup>+</sup> | Roche-Merck | 10127965001 |
| RNase Inhibitor, Murine | New England Biolabs | M0314 |
| KLD Enzyme Mix | New England Biolabs | E0554 |
| Gateway BP Clonase | Thermo Fisher Scientific | 11789020 |
| Gateway LR Clonase | Thermo Fisher Scientific | 11791020 |
| E. coli Poly(A) Polymerase | New England Biolabs | M0276 |
| Proteinase K | New England Biolabs | P8107 |
| Thermolabile Proteinase K | New England Biolabs | P8111 |
| Monarch RNA Cleanup Kit | New England Biolabs | T2030 |
| TALON Metal Affinity Resin | Takara Bio | 635502 |
| SYBR<sup>TM</sup> Gold Nucleic Acid Gel Stain | Thermo Fisher Scientific | S11494 |
| Benzonase Nuclease | Merck Millipore | 70746 |
| Gaussia Luciferase Assay (GAR-2B) | Targeting Systems | GAR-2B |
| Phusion High-Fidelity DNA Polymerase | New England Biolabs | M0530 |
| **Critical commercial assays** |  |  |
| PURExpress In Vitro Protein Synthesis Kit | New England Biolabs | E6800 |
| AMP-Glo Assay | Promega | V5011 |
| MEGAscript T7 Transcription Kit | Invitrogen | AM1334 |
| **Deposited data** |  |  |
| N/A |  |  |
| **Experimental models: Cell lines** |  |  |
| HEK293T | ATCC | RRID: CVCL_0063 |
| HeLa-S3 | ATCC | RRID: CVCL_0058 |
| U2OS | ATCC | RRID: CVCL_0042 |
| HMC3 | ATCC | RRID: CVCL_II76 |
| LN-229 | ATCC | RRID: CVCL_0393 |
| A-172 | ATCC | RRID: CVCL_0131 |
| SH-SY5Y | ATCC | RRID: CVCL_0019 |
| Experimental models: Organisms/strains |  |  |
| N/A |  |  |
| DNA template used for in vitro transcription, DNA gBLOCKs, DNA primers used for cloning, site-directed mutagenesis and sgRNA are given in Supp. Table S2 |  |  |
| Recombinant DNA |  |  |
| pDONR221-APTX | This study | Addgene *Will follow later* |
| pDONR221-ScARP | This study | Addgene *Will follow later* |
| pDONR221-TARG1 | <sup>26</sup> | Addgene 172577 |
| pDONR221-PARP15cat | This study | Addgene *Will follow later* |
| pDONR221-PARP15catHY | This study | Addgene *Will follow later* |
| pDONR221-NUDT5 | <sup>12</sup> | Addgene 219954 |
| pDEST17-APTX | This study | Addgene *Will follow later* |
| pDEST17-ScARP | This study | Addgene *Will follow later* |
| pDEST17-TARG1 | <sup>26</sup> | Addgene 172594 |
| pDEST17-TRPT1 | <sup>9</sup> | Addgene 184239 |
| pDEST17-PARP15cat | Addgene | Addgene *Will follow later* |
| pDEST17-NUDT5 | <sup>12</sup> | Addgene 219955 |
| Software and algorithms |  |  |
| R (version 4.4.2) | R Foundation | https://www.r-project.org |
| ggplot2 | Wickham et al., 2016 | https://ggplot2.tidyverse.org |
| ImageJ / FIJI | NIH | https://imagej.nih.gov/ij |
| Inkscape 1.4 | Inkscape Project, 2020 | https://inkscape.org |
| PyMol v.2.1 | Schrödinger | https://pymol.org/2/ |

### EXPERIMENTAL MODEL AND STUDY PARTICIPANT DETAILS

All E. coli strains used in this study were grown in Luria-Bertani (LB) broth (Fisher Scientific) supplemented with 25 µg/mL chloramphenicol, 50 µg/mL kanamycin or 100 µg/ml ampicilin. HEK293T and HeLa cells were obtained from American tissue culture collection (ATCC), routinely tested for mycoplasma contamination and used at low passage (<20 passages) after thawing.

### METHOD DETAILS

#### Materials, reagents and chemicals

High-fidelity DNA polymerase Phusion, KLD enzyme mix and Gateway cloning reagents were obtained from New England Biolabs and Thermo Scientific. All the DNA and RNA oligonucleotides used in the study were synthesized by Integrated DNA Technologies and sequences are given in Table S1. The oligonucleotides were resuspended to 100 µM stock solutions in 20 mM HEPES–KOH (pH 7.6), 50 mM KCl buffer or RNase free water. Source of other chemicals used in the study is provided in Table S1

#### Mammalian and bacterial expression constructs

The generation of the mEGFP-PARP15 expression plasmid was described previously^27^ and is available from Addgene. The H559Y catalytically inactive mutant of PARP15 was generated by site-directed mutagenesis using Phusion High-Fidelity DNA Polymerase and the KLD Enzyme Mix (New England Biolabs). Primers used for mutagenesis are listed in Table S1. The construction of TRPT1 bacterial expression plasmids has been described previously^9^. The human PARG (residues 448–976) cloned into the pNH-TrxT vector was a kind gift from Lari Lehtiö ^28^. Cloning of additional hydrolases was performed as described previously^29,30^. The full-length human ARH3 was a gift from Bernhard Lüscher and was PCR-cloned into the pDONR221-Kan entry vector and subsequently transferred into the pDEST17 destination vector using Gateway cloning for bacterial expression. For APTX and ScARP, a gBlock fragment (IDT) was synthesized to enable insertion into pDONR221, and the construct was then subcloned into pDEST17 using Gateway technology. The sequence of the APTX and ScARP gBlock is provided in Table S1, along with Addgene accession numbers for all plasmids deposited. Single guide RNAs (sgRNAs) targeting the human TARG1 locus were cloned into the Cas9/puromycin expression vector pSpCas9(BB)-2A-Puro V2.0 using BbsI restriction sites as described earlier^31^. Complementary oligonucleotides annealed, and phosphorylated and the resulting duplex was ligated into BbsI-digested pX459. Ligation mixtures were transformed into chemically competent Escherichia coli, and plasmids from individual colonies were verified by Sanger sequencing to confirm correct sgRNA insertion.

#### Recombinant proteins expression and purification

PARP10cat and TRPT1 were purified as described earlier^9^, purification of hydrolases was also described in detail before^12^. Recombinant human APTX was expressed in E. coli BL21(DE3) cells carrying the pDEST17-APTX plasmid. A 50 mL LB pre-culture (Amp^R) was grown overnight at 37°C and used to inoculate 2 L of LB medium. Cultures were grown at 37°C until reaching an OD_600_ of 0.6–0.8, then induced with 0.4 mM IPTG and 2% ethanol. Protein expression was carried out overnight at 18°C. ScARP expression in E. coli BL21(DE3) was induced with 0.8 mM IPTG at 30°C for 3 hours as described^32^. Cells were harvested by centrifugation at 6,000 rpm for 15 min at 4°C and stored at -80°C. Cell pellets were resuspended in 20 mL/g lysis buffer (50 mM Tris-HCl pH 7.5, 500 mM NaCl, 5% glycerol, 0.5 mM TCEP, 10 mM imidazole, 0.1% NP-40, protease inhibitor cocktail [EDTA-free], and Benzonase), supplemented with 500 µg/mL lysozyme. After 30 min incubation on ice, cells were sonicated (3 min total, 30 s on / 40 s off, 20% amplitude) and centrifuged at 12,000 g for 60 min at 4°C. The soluble fraction was incubated with 0.6 mL pre-equilibrated 3P TALON metal affinity resin (Clontech) for 1 h at 4°C while rotating. After binding, the resin was washed four times with wash buffer (50 mM Tris-HCl pH 7.5, 200 mM NaCl, 10 mM imidazole), and protein was eluted with three fractions of 0.4 mL elution buffer (50 mM Tris-HCl pH 7.5, 500 mM NaCl, 300 mM imidazole). Elutions were dialyzed overnight at 4°C in 4 L of dialysis buffer (50 mM Tris-HCl pH 7.5, 150 mM NaCl). His tagged catalytic domain of PARP15 was purified essentially the same protocol additional high salt (1M NaCl) wash was included during purification to remove potentially associated RNases. NADAR was purified using the same protocol but Rosetta cells were used for expression. After dialysis proteins were supplemented with 20% glycerol, snap-frozen in liquid nitrogen and stored at -80°C.

#### ADP-ribosyltransferase and hydrolase assays

ADP-ribosylation and hydrolase reactions were performed in ADPr reaction buffer (20 mM HEPES–KOH pH 7.6, 50 mM KCl, 5 mM MgCl_2_, 1 mM DTT, 500 µM *β*-NAD^+^, and 40 U RNase inhibitor [New England Biolabs]) at 37°C for the indicated times. RNA oligonucleotide substrates (typically 1 µM) were incubated with 1-2 µM of the respective transferase or hydrolase, unless stated otherwise. Following the reaction, proteins were digested with 20 U Proteinase K (New England Biolabs) for 20 min at 37°C, and samples were resolved on denaturing urea-PAGE. For ADP-ribosylhydrolase assays, RNA oligonucleotides were first modified in 100-200 µL ADP-ribosylation reactions, then purified using the Monarch® Spin RNA Cleanup Kit (New England Biolabs) before use in hydrolase reactions. After completion of the reactions, samples were mixed with RNA loading dye containing urea and denatured for 3 min at 95°C. Samples were then loaded onto pre-run urea-polyacrylamide gels (8 M urea, 20% acrylamide:bisacrylamide [19:1], 0.2% APS, 0.4% TEMED). Gels were run in TBE buffer (89 mM Tris, 89 mM boric acid, 2 mM EDTA, pH 8.0) at a constant 7 W. Where indicated, boronate-containing urea-PAGE gels were prepared by supplementing with 0.25% 3-acrylamidophenylboronic acid.

#### LC-MS/MS

Degradation of in vitro ADP-ribosylated RNA as well as cellular RNA was performed using a procedure adopted from (PMID: 26751644). LC-MS/MS analysis was performed on an Agilent 1290 Infinity Binary LC system (Agilent Technologies) using ZORBAX SB-C18 column (Agilent Technologies, 2.1 mm × 50 mm, 1.8 µm) coupled to an Agilent 6490 triple quadrupole mass spectrometer. Elution was performed with 5 mM ammonium acetate pH 6.9 and acetonitrile (ACN), the flow was 0.3 ml/min from 0-7 min, then linearly increased from 0.3 ml/min to 0.38 ml/min in 7-10.5 min, then switched to 0.5 ml/min for 10.5 - 14.5 min, and 0.3 ml/min for 14.5 - 15.0 min. The column was kept at 30 °C. The gradient was: 0 - 3 min, 0% ACN; 3 - 7 min, 0 - 5% ACN; 7 - 10.5 min, 5 % ACN, 10.5 -12.5 min 5 - 50% ACN; 12.5 - 15.0 min, 0% ACN. The MS source-dependent parameters were as follows: gas temperature 110 °C, gas flow 19 l/min (N2), nebulizer 25 psi, sheath gas heater 375 °C, sheath gas flow 11 l/min (N2), capillary voltage 2000 V (positive mode), nozzle voltage 0 V, fragmentor voltage 300 V, high pressure RF 70 V and low pressure RF 80 V. Compound dependent parameters are listed in Table S1.

#### Cell culture and transfection

All cells were kept at a humidified atmosphere at 37°C with 5% CO2 and were cultivated in DMEM with high glucose and GlutaMAX™ (Gibco) supplemented with 10% heat-inactivated foetal bovine serum (Gibco). HEK293T cells were seeded in 6-well microplates with 50% confluency and transfected next day using calcium phosphate, for which a detailed protocol is available ^33^. The following day, cells were washed with warm HEPES, followed by RNA or protein extraction 48h after transfection. For subsequent analysis of ADPr-RNA using LC-MS/MS similar approaches were followed, except that cells were grown in 10cm dishes. TARG1 knock-out cells were made using CRISPr-Cas9 method. Cells were transfected as described with pX459-sgTARG1 plasmid and then selected with 1 µg/ml of puromycin. After 24h cells were diluted in a 150 cm dish and individual colonies picked using cloning disks after 1-2 weeks. TARG1 knock-out was verified using a TARG1 specific antibody and western blot. To make inducible TARG1 knock-down cells, TARG1 shRNA oligos were annealed and cloned into AgeI-EcoRI site of Tet-plko-Puro plasmid (Addgene 21915). After confirming the insertion with a restriction digest, the plasmid was co-transfected with pCMV-VSVG and and pCMV-dr8.2 dvpr into HEK293T cells using calcium-phosphate transfection. Medium with viral particles was harvested after 24h, fresh medium added and then viral particles harvested after another 24 h incubation. The two viral supernatants were pooled, centrifuged 500xg for 5 min and filtered through a 0.45 μm PES filter. Target cells were transduced in the presence of polybrene with 1-2 ml of medium containing viral particles per 10 cm dish. After 24 h fresh medium with puromycin (1-3 µg/ml) was added and cells cultivated for 1-2 weeks.

#### Western blot

Cell protein extractions were performed using RIPA buffer (150 mM NaCl, 1% Triton X-100, 0.5% sodium deoxycholate, 0.1% SDS, 50 mM Tris–HCl (pH 8.0)) supplemented with protease inhibitor cocktail, followed by benzonase treatment (Santa Cruz Biotechnology). Proteins were separated on 12% gels and blotted onto nitrocellulose. Membranes were blocked with 5% non-fat milk in PBST (PBS with 0.05% Tween 20) for 1 hour at RT, primary antibodies were diluted in PBST and incubated overnight in a cold room, secondary antibodies were diluted in 5% non-fat milk in PBST and incubated for 1 h at RT. Wash steps were performed in between and after antibody incubations with PBST at RT for at least 5 min. The HRP-conjugated Peroxidase AffiniPure Goat Anti-Rabbit IgG (H + L) secondary antibody (Jackson) was diluted 1:10.000 in 5% non-fat milk in PBST and incubated for 1 hour at RT. Chemiluminescence signals were detected using the Azure600. Transient overexpression of mEGFP-PARP15 was detected using a mouse polyclonal anti-GFP antibody (600-101-215; Rockland). TARG1 knockout and knockdown were confirmed with TARG1 antibody 3A5^26^ (diluted 1:200).

#### RNA isolation and detection of ADPr-RNA using AMPGlo assay

Total RNA was isolated using TRIzol (Thermo Fisher Scientific) according to the manufacturer’s instructions with minor modifications. Briefly, cells were lysed directly in 1 ml of TRIzol reagent per 10 cm dish, and phase separation was induced by addition of chloroform followed by centrifugation. The aqueous phase was transferred to a fresh tube, and RNA was precipitated with cold isopropanol, washed twice with 75% ethanol, and resuspended in nuclease-free water. RNA quantity and purity were assessed by spectrophotometry (A260/280), and integrity was verified by agarose gel electrophoresis. For the determination of hydrolase activity using AMPGlo, modified RNA oligos were purified after transferase reaction using Monarch RNA Cleanup Kit. The indicated amounts of ADPr modified oligonucleotide or isolated cellular RNA were incubated with indicated hydrolases in AMPGlo buffer (50 mM Tris, pH 7.0, 100 mM NaCl, 5 mM MgCl_2_) and incubated for 30 min at 37°C and then additional 30 min with 200 nM NUDT5. AMP was measure using AMP-Glo™ assay (Promega) performed with 10 µl reaction in 384 well plates according to the manufacturers protocol. Luminescence was measured using a SpectraMax iD3.

#### TARG1 RNA binding assay

RNA electrophoretic mobility shift assays (EMSAs) were performed essentially as described^**30**^. pcDNA3.1-based Pentaprobe plasmids were linearized with ApaI and 3′ overhangs were filled in using Klenow polymerase. One microgram of purified, linearized DNA was transcribed in vitro with the T7 RiboMAX Large Scale RNA Production System (Promega) in the presence of α-^32^P-UTP. Transcription reactions were resolved on denaturing urea-TBE polyacrylamide gels, and the gel slice containing the labeled Pentaprobe RNA was excised, crushed, and eluted in water overnight. After centrifugation, the supernatant was collected, the gel pellet was re-soaked for 2 h, and combined eluates were ethanol-precipitated overnight at -20°C. Pellets were centrifuged (30 min, 4°C, 13,000 × g), air-dried at 37°C, resuspended in 100-150 µl TBE buffer, incubated for 5-10 min at 37°C, and stored at -20°C. Immediately before use, probes were denatured for 45 s at 95°C and snap-cooled on ice. RNA binding reactions (30 µl) contained 1 µl of radiolabeled RNA and 0-8 pmol recombinant protein in the gel-shift buffer (10 mM MOPS [pH 7.0], 50 mM KCl, 5 mM MgCl_2_, 1 mM DTT, 10% glycerol) and were incubated for 30 min at 4°C. Complexes were resolved on 7% native polyacrylamide (acrylamide:bisacrylamide 19:1) gels in TB buffer (45 mM Tris, 45 mM boric acid) at 10 mA and 4°C for 2-3 h. Gels were dried and analyzed by autoradiography.

#### In vitro translation assay

gBLOCK of codon optimised Gluc was synthesized by IDT. 5’UTR contained T7 promoter and ribosomal binding site and 3’UTR contained stop codon and bacterial terminator hairpin to enable efficient translation termination. After PCR amplification GLuc reporter was generated by *in vitro* transcription using 1 *µ*g of template DNA and incubation at 37°C for 4 h using MEGAscript™ T7 Transcription Kit (Invitrogen). GLuc reporter was poly(A) tailed using E. coli Poly(A) Polymerase (New England Biolabs). IVT reactions were DNase treated for 30 min at 37°C, using 1 µl TURBO DNase (Invitrogen). RNA was purified using the Monarch RNA Cleanup Kit (New England Biolabs). Produced RNA was then ADP-ribosylated using recombinant PARP15cat. After proteinase K treatment and purification ADPr-RNA Gluc was enriched using Anti-PAR/MAR (E6F6A0, Cell Signalling Technology (CST)) coupled to protein A/G magnetic agarose beads (Pierce). After elution ADPr-Gluc or non-modified Gluc mRNA were used *in vitro* translation assays using the PURExpress in vitro protein synthesis kit (New England Biolabs) according to the manufacturer’s protocol. GLuc activity was measured with the GAR-2B Gaussia Luciferase assay (Targeting Systems).

### QUANTIFICATION AND STATISTICAL ANALYSIS

All experiments were replicated at least once, with results similar to the data displayed, and no data were excluded from analysis. Urea-PAGE signals were quantified using ImageJ. Luminescence signals from AMP-Glo assays were analysed in R; the base R package was used for statistical testing and ggplot2 for data visualization. Released AMP was quantified by generating a calibration curve from an AMP standard series measured in relative light units (RLU), subtracting the mean blank signal, and fitting a linear regression to log-transformed RLU versus log-transformed AMP concentration to obtain calibration coefficients. Background-corrected RLU values from experimental samples were converted to AMP concentrations (µM) using this calibration model and then to total picomoles per well based on the reaction volume; where indicated, these values were further normalized to input RNA and expressed as pmol of AMP per microgram RNA. Unless stated otherwise, data are presented as mean ± standard error of the mean (SEM) of independent biological replicates (n), where n denotes independently transfected and processed cell cultures. Unpaired Welch’s t-tests were used where appropriate to compare groups; exact P values are reported in the figures. No formal randomization or a priori sample-size estimation was applied; all successfully measured biological replicates were included, and data distribution and variance were inspected visually (dot plots and residuals) to confirm that the assumptions of the parametric tests were reasonably met.

