## Supplementary figures and tables for "Human PARPs modify RNA nucleobases *in vitro* and in cells"

### SUPPLEMENTAL INFORMATION

### SUPLEMENTARY FIGURES

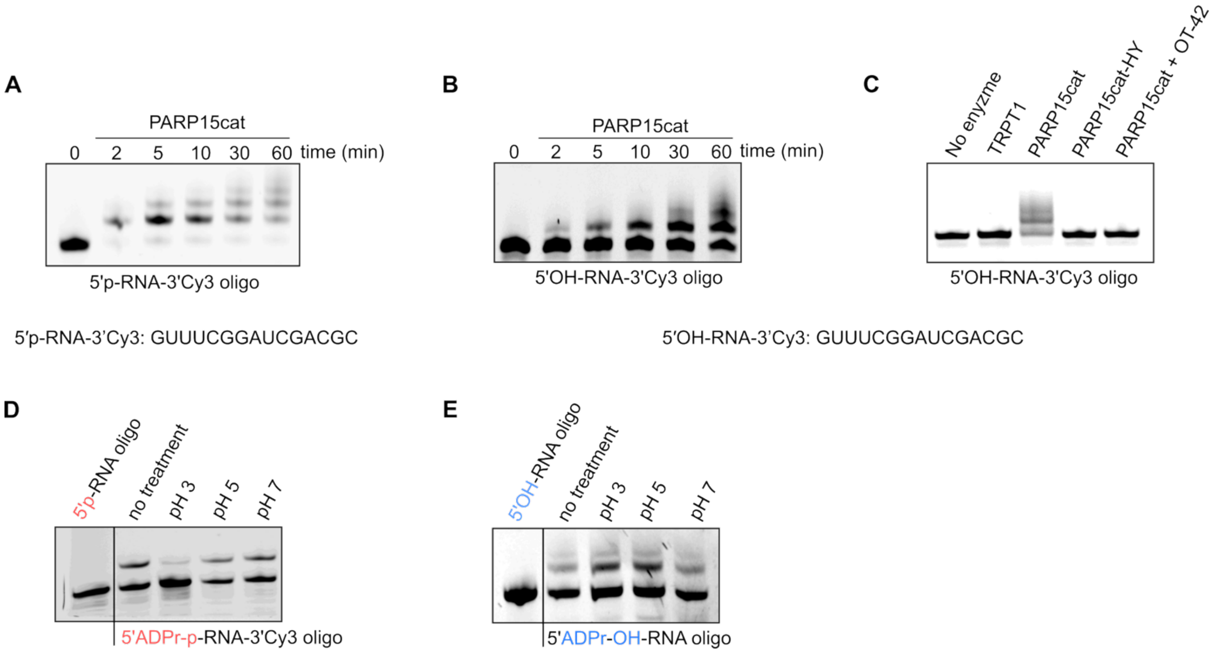


**Supplementary Figure S1. ADP-ribosylation activity of PARP15cat and TRPT1, related to Figure 1. (A)** Time dependent *in vitro* ADP-ribosylation activity of PARP15cat on 5' monophosphorylated RNA oligo (5'p-RNA-3'Cy3 oligo). **(B)** Time dependent in vitro ADP-ribosylation activity of PARP15cat on 5' hydroxyl RNA oligo (5'OH-RNA-3'Cy3 oligo). **(C)** In vitro ADP-ribosylation activity of PARP15cat and catalytically inactive mutant H559Y and inhibition of PARP15cat by PARP10/15 inhibitor OT-42. Inhibition of PARP15cat with OT-42. The 5' hydroxyl RNA oligo (5'OH-RNA-3'Cy3 oligo) was used as substrate. All gels are representative of two to three independent experiments. **(D-E)** A purified 5'-ADPr-p-oligo modified with TRPT1 (D) or 5'-ADPr-OH-oligo modified with PARP15cat (E) was incubated in buffers with different pH values as indicated and analysed using urea-PAGE and in-gel fluorescence detection.

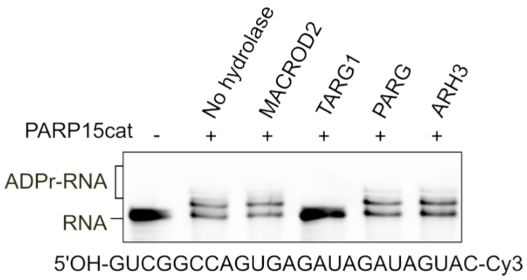


**Supplementary Figure S2. Hydrolase activity of different hydrolases ADPr-RNA oligo related to Figure 3.** ADPr-RNA oligo containing all four different ribonucleotides was ADP-ribosylated *in vitro* by PARP15cat and purified after proteinase K digest. Activity of different hydrolases was assessed by incubating ADPr-RNA oligo with 1 µM of indicated hydrolases for 30 min at 37ºC. Reactions were resolved on 20% urea-PAGE and visualized using in-gel fluorescence.

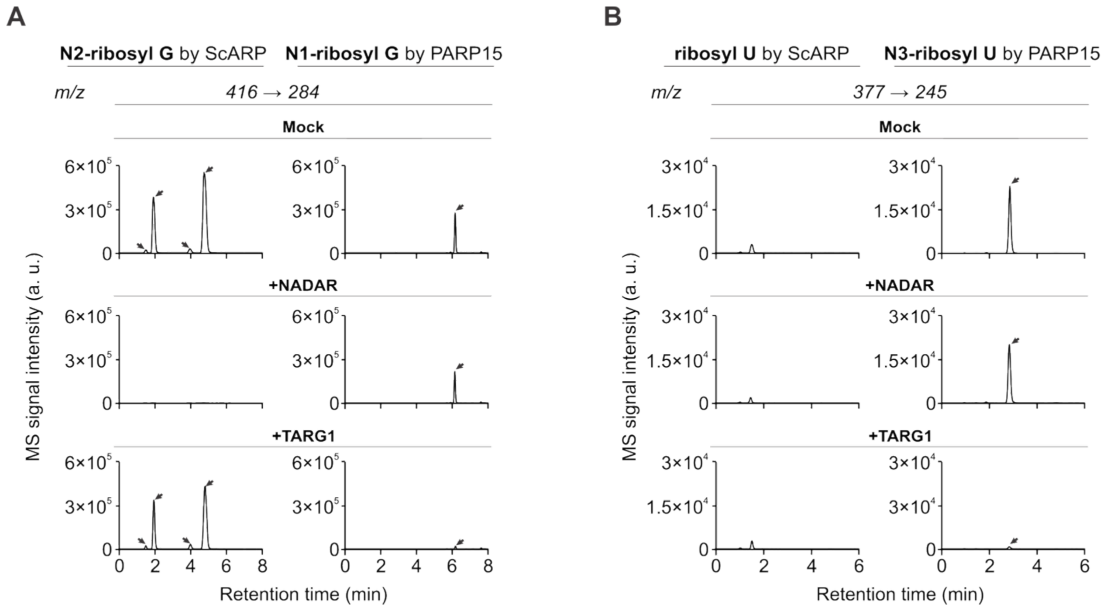


**Supplementary Figure S3. LC-MS/MS analysis of reaction products generated by ScARP and PARP15 activity related to Figure 3.** LC-MS/MS chromatograms of reaction products from in vitro ADP-ribosylation assays in the presence of PARP15 or ScARP and treated with NADAR, TARG1 or non-treated (Mock). (A) LC-MS/MS chromatograms guanine-ADPr and (B) uracil-ADPr.

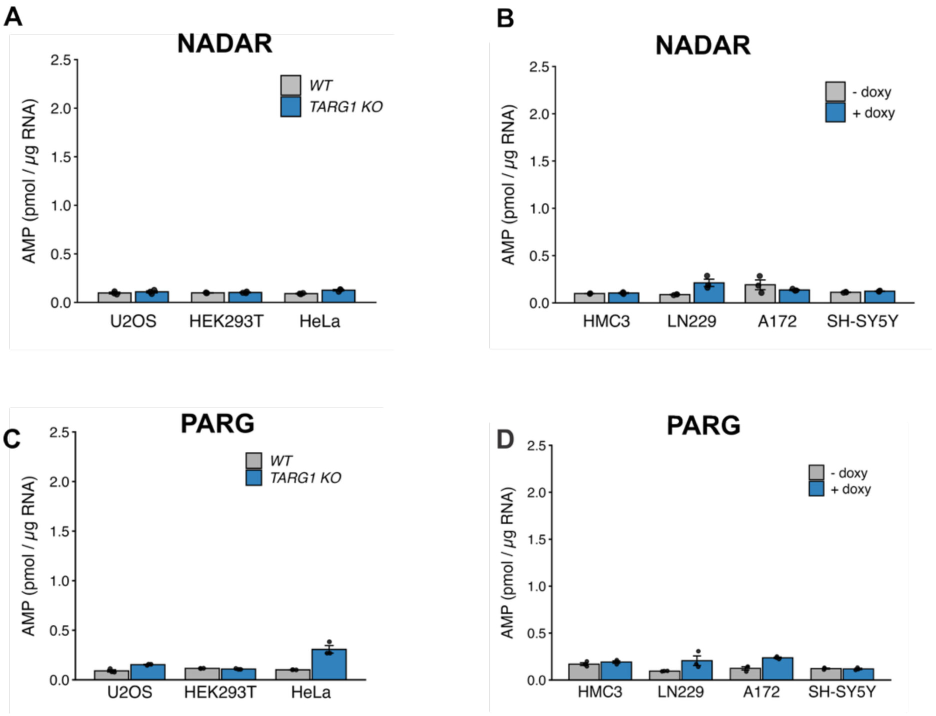


**Supplementary Figure S4. ADPr release measured by AMP-Glo assay related to Figure 4. (A)** RNA was isolated from U2Os, HEK293T or HeLa TARG1 knockout cells. Extracted RNA was first incubated with 100 nM NADAR followed by NUDT5 and measurement of AMP levels using AMP-Glo. (B) HMC3, Ln229, A172 and SH-SY5Y were kept with or without doxycycline for 72 h prior to RNA extraction. Extracted RNA was first incubated with 100 nM NADAR followed by NUDT5 and measurement of AMP levels using AMP-Glo. (**C**) As in (A), but with incubation of RNA with PARG prior to AMP-Glo. (**D**) As in (B), but with incubation of RNA with PARG prior to AMP-Glo.

### SUPLEMENTARY TABLES

#### Supplementary Table S1. MRM transitions used in this study.

| Nucleoside | Precursor ion (*m/z*) | Fragment ion (*m/z*) | Collision energy | Cell accelerator voltage |
|---|---|---|---|---|
| ribosyl-U | 377 | 245 | 2 | 6 |
| $^{13}C_9$-ribosyl-U | 386 | 254 | 2 | 6 |
| r-rG | 416 | 284 | 7 | 4 |
| $^{13}C_{10}$-ribosyl-G | 426 | 294 | 7 | 4 |
| ‘nucleoside 387’ | 387 | 255 | 2 | 6 |
| ‘nucleoside 425’ | 425 | 293 | 7 | 4 |

#### Supplementary Table S3. Oligonucleotides, gBLOCKS and constructs used in this study, related to STAR methods.

| Oligo name / modifications | Sequence (5’->3’) | Purpose |
|---|---|---|
| 5’OH-poly(U)-3’Cy3 | UUUUUUUUUUUUUUU | ADP-ribosylation assay |
| 5′p-RNA-3’Cy3 | GUUUCGGAUCGACGC | ADP-ribosylation assay |
| 5′OH-RNA-3’Cy3 | GUUUCGGAUCGACGC | ADP-ribosylation assay |
| 5’OH-poly(AG)-3’Cy3 | AGAGAGAGAGAGAGAG | ADP-ribosylation assay |
| AO-035 (5′OH-RNA-3’) | GUUUCGGAUCGACGC | LC-MS/MS (Figure 2B) |
| AO-036 (DNA duplex) | CAGTAATACGACTCACTATTAGTTGGTGGTTGTTGTGTG<br>TTTGTGGTTGGTTTGTTTGG | *in vitro* transcription and LC-MS/MS (Figure 2C, 2D, 2F, 2G) |
| sgTARG1-f | CACCGATCAGTGAGGATTGTCGCATGGG | Used for making TARG1 knock-out cells |
| sgTARG1-r | AAACCCCATGCGACAATCCTCACTGATC | |
| shTARG1-f | CCGGT GAGAGATGGGCGATATATA CTCGAG<br>TATATATCGCCCATCTCTC TTTTTG | Used for making TARG1 knock-down cells |
| shTARG1-r | AATTCAAAAA GAGAGATGGGCGATATATA CTCGAG<br>TATATATCGCCCATCTCTC A | |
| Pentaprobes | 5’-TAAAGTAAGA TAAGGCAAGA CAAGGTAAAA<br>CGGAACAGAA CCGAAGGGAA GAGAAGCAAA<br>GCGAAAGGAA ATAAACAAAA<br>GAAAAATTCG TAGAATTCCG-3’ (PP7) | Used for EMSA [1] |
| His-NUDT5 | ATGTCGTACTACCATCACCATCACCATCACTCGGATGAA<br>AACCTGTATTTTCAGGGCAGACTCGAGGGTGGTGGCGG<br>TTCAGGCGGAGGTGGCTCTCTTGAAGTCCTCTTTCAGG<br>GACCCACAAGTTTGTACAAAAAAGCAGGCTTCGAGAGC<br>CAAGAGCCAACCGAAAGCTCCCAGAACGGGAAACAGTA<br>CATTATTTCGGAAGAGTTAATCAGCGAAGGTAAGTGGGT<br>CAAACTTGAAAAGACCACCTACATGGATCCTACTGGCAA<br>GACACGTACATGGGAATCTGTTAAACGTACTACACGTAA<br>GGAACAGACCGCTGATGGAGTTGCCGTAATTCCAGTTC<br>TTCAGCGTACTCTTCACTACGAATGCATCGTTCTGGTAA<br>AGCAGTTTCGCCCACCGATGGGAGGTTATTGTATTGAG<br>TTTCCCGCTGGTCTTATCGATGACGGTGAGACACCAGA<br>GGCTGCGGCATTGCGTGAGCTTGAGGAAGAAACCGGTT<br>ATAAGGGCGACATCGCGGAGTGTTCACCAGCTGTTTGT<br>ATGGACCCGGGTTTATCAAACTGTACTATTCACATTGTA<br>ACTGTCACGATCAATGGTGACGATGCAGAGAACGCTCG<br>TCCAAAACCGAAACCTGGCGATGGCGAGTTTGTAGAGG<br>TAATCTCCTTGCCAAAGAACGACCTTTTACAGCGCTTGG<br>ACGCCCTTGTTGCTGAGGAACATTTGACAGTTGATGCG<br>CGTGTTTACTCTTACGCCTTGGCGCTTAAACACGCCAAC<br>GCTAAGCCGTTTGAAGTACCCTTCTTAAAGTTTTAA | Purification from E. coli |
| His-PARP15cat | ATGTCGTACTACCATCACCATCACCATCACCTCGAATCA<br>ACAAGTTTGTACAAAAAAGCAGGCTCGATGAATCTTCCT<br>GAACACTGGACTGACATGAATCATCAGCTGTTTTGCATG<br>GTCCAGCTAGAGCCAGGACAATCAGAATATAATACCATA<br>AAGGACAAGTTCACCCGAACTTGTTCTTCCTACGCAATA | Purification from E. coli |

| | | |
|---|---|---|
| | GAGAAGATTGAGAGGATACAGAATGCATTTCTCTGGCA GAGCTACCAGGTAAAGAAAAGGCAAATGGATATCAAGA ATGACCATAAGAATAATGAGAGACTCCTCTTCCATGGGA CAGATGCAGACTCAGTGCCATATGTCAATCAGCACGGC TTTAATAGAAGTTGTGCTGGGAAAAATGCTGTATCCTAT GGAAAAGGAACCTATTTTGCTGTGGATGCCAGTTATTCT GCCAAGGACACCTACTCCAAGCCAGACAGCAATGGGAG AAAGCACATGTACGTTGTGCGAGTACTTACTGGAGTCTT CACAAAGGGACGTGCAGGATTAGTCACCCCTCCACCCA AGAATCCTCACAATCCCACAGATCTCTTTGACTCAGTGA CAAACAATACACGATCTCCAAAGCTATTTGTGGTATTCT TTGATAATCAGGCTTACCCAGAATATCTCATAACTTTCAC GGCTTGA | |
| His-NADAR | ATGTCGTACTACCATCACCATCACCATCACCTCGAATCA ACAAGTTTGTACAAAAAAGCAGGCTTCGCAGTACGCCC CGTCTTCGTTCCGACAAATGCGGGTAACCTGTTATCAAT CACGAAAGACGTGGACTTCCCGTGGGCACCCGGCATG AGTAAAACGCAAAAACAAAAGTCTATTCGCGCCCTGCAC ACGGCGGCCAACGAACAGGGCCTTAACTCTTTACTTGA AATCTCGAGCAAATCCGAAGATGCACTGGGAGTCGCCC TGAGTGCTTTCAATCTTCGTATCAAAACTAAGCGTTTAG GAAAGGAATTCACTGTTGAAAGCGCGTTCCAGGCAAGC AAAGTTTTTGAAATGGGTGGACCCTATGTCGATATTTTA GACAAATCGTCCATCGAGGCGAAGAAAGATATGCGCTT GAAAGAGTCGGGGGGATTAGTAAATTTCAAGTTTTATAA CACCATTTGGCCTATTGTACCTCGCACGGCATTCTATGA CTGGTTGTATCTGTCCGCGCTGAATCAGAACAAGAACC TTGCTTTGCACTTATTGAATTTCGACGGATTCACCGACA TTGAATTCAATCCCGCTAAAAGTATCAATTGCCAGGCCC GTGCCGCTGCTCTTTTTGTATCGCTGGTACGTCGCAACA TGCTGGATGATGTGTTGAGCAGCAAGGACGGTTTCCTG TCGAAATTGGCGTCGCATTACGGGGTGGAAAACTACAG TATCCAGCACACCTTAATAtga | Purification from E. coli |
| His-ScARP | CATCACCATCACCATCACCTCGAATCAACAAGTTTGTAC AAAAAAGCAGGCTTCatgCCGTCGGCTGCCCCCGCCAAG GCCGCCCCGGCCTGCCCCCAGTTCGACGACCGGACCA AGGCCGCCGCCGACCGCGGCGTCGACGTCGACCGCAT CACGCCCGAGCCGGTCTGGCGCACCACCTGCGGCACC CTCTACCGCAGCGACAGCCGCGGCCCGCAGGTCGTCT TCGAGGAGGGCTTCCACGCCAAGGACGTCCAGAACGG ACAGTACGACGTCGAGAAGTACGTCCTGGTCAACCAGC CCTCGCCGTACGTGTCGACGAGCTACGACCACGACCTG TACAAGACCTGGTACAAGTCCGGCTACAACTACTACGT CGACGCCCCCGGCGGCATCGACGTCAACAAGACCATC GGCGACACCCACAAGTGGGCCGACCAGGTCGAGGTCG CCTTCCCGGGCGGCATCCAGCGGAAGTACATCATCGG CGTCTGTCCGGTCGACCGGCAGACCAAGACCGAGATC ATGAGCGACTGCGAGAGCAACCCGCACTACCAGCCCT GGCACTGA | Purification from E. coli |
| Gluc-PE | TAATACGACTCACTATAGGGAAGGAGGTTTAAATATGAA ACCGACCGAAAATAATGAAGATTTTAATATTGTGGCGGT GGCGAGCAATTTTGCGACCACCGATCTGGATGCGGATC GTGGCAAACTGCCGGGCAAAAAACTGCCGCTGGAAGT GCTGAAAGAAATGGAAGCGAATGCGCGTAAAGCGGGCT GCACCCGTGGCTGCCTGATTTGCCTGAGCCATATTAAAT GCACCCCGAAAATGAAAAAATTTATTCCGGGCCGTTGC CATACCTATGAAGGCGATAAAGAAAGCGCGCAAGGCGG CATTGGCGAAGCGATTGTGGATATTCCGGAAATTCCGG GCTTTAAAGATCTGGAACCGATGGAACAATTTATTGCGC AAGTGGATCTGTGCGTGGATTGCACCACCGGCTGCCTG AAAGGCCTGGCGAATGTGCAATGCAGCGATCTGCTGAA AAAATGGCTGCCGCAACGTTGCGCGACCTTTGCGAGCA AAATTCAAGGCCAAGTGGATAAAATTAAAGGCGCGGGC GGCGATtaataaGGATACAATGTTTCGAAACATTGTATCC | *In vitro* translation (Figure 4J) |
| Gluc | CAGTAATACGACTCACTATAGGGAGGACTCACTATTTGT TTTCGCGCCCAGTTGCAAAAAGTGTCGCCACCATGGGA GTCAAAGTTCTGTTTGCCCTGATCTGCATCGCTGTGGCC GAGGCCAAGCCCACCGAGAACAACGAAGACTTCAACAT CGTGGCCGTGGCCAGCAACTTCGCGACCACGGATCTC | In cells translation (Figure 4I) |

| | | |
|---|---|---|
| | GATGCTGACCGCGGGAAGTTGCCCGGCAAGAAGCTGC<br>CGCTGGAGGTGCTCAAAGAGATGGAAGCCAATGCCCG<br>GAAAGCTGGCTGCACCAGGGGCTGTCTGATCTGCCTGT<br>CCCACATCAAGTGCACGCCCAAGATGAAGAAGTTCATC<br>CCAGGACGCTGCCACACCTACGAAGGCGACAAAGAGT<br>CCGCACAGGGCGGCATAGGCGAGGCGATCGTCGACAT<br>TCCTGAGATTCCTGGGTTCAAGGACTTGGAGCCCATGG<br>AGCAGTTCATCGCACAGGTCGATCTGTGTGTGGACTGC<br>ACAACTGGCTGCCTCAAAGGGCTTGCCAACGTGCAGTG<br>TTCTGACCTGCTCAAGAAGTGGCTGCCGCAACGCTGTG<br>CGACCTTTGCCAGCAAGATCCAGGGCCAGGTGGACAA<br>GATCAAGGGGGCCGGTGGTGACTAAGAGAGCTCGCTTT<br>CTTGCTG | |
| His-APTX | ATGTCGTACTACCATCACCATCACCATCACCTCGAATCA<br>ACAAGTTTGTACAAAAAAGCAGGCTTCATGATGCGTGTA<br>TGCTGGCTTGTCCGTCAGGATTCACGTCATCAGCGTATT<br>CGTCTGCCTCACTTAGAGGCGGTTGTCATTGGTCGTGG<br>ACCAGAGACTAAAATTACCGATAAAAAGTGCAGTCGCCA<br>ACAAGTTCAGCTGAAAGCGGAATGCAACAAGGGATATG<br>TTAAGGTCAAGCAAGTCGGAGTCAATCCTACGTCTATTG<br>ATTCCGTTGTCATCGGGAAGGACCAAGAGGTCAAGTTA<br>CAACCGGGTCAGGTTCTGCACATGGTCAATGAGCTTTA<br>CCCATACATTGTAGAATTTGAGGAGGAAGCGAAAAATCC<br>TGGGTTGGAAACACACCGCAAGCGTAAGCGTTCGGGTA<br>ACTCGGACAGCATTGAACGTGACGCGGCCCAAGAAGCA<br>GAGGCAGGTACTGGTCTTGAACCCGGCAGCAACTCCG<br>GGCAATGTTCGGTGCCATTAAAAAAGGGTAAGGACGCA<br>CCGATCAAGAAGGAATCTTTGGGCCATTGGTCACAGGG<br>CTTGAAGATCAGTATGCAGGACCCTAAAATGCAAGTCTA<br>TAAGGACGAACAGGTTGTCGTCATCAAAGATAAGTATCC<br>CAAAGCCCGTTACCACTGGCTTGTGCTGCCCTGGACAT<br>CGATCAGCTCATTGAAAGCTGTTGCGCGCGAACATCTG<br>GAGTTGTTGAAACATATGCATACTGTGGGAGAAAAGGT<br>CATCGTTGACTTTGCCGGGTCCTCGAAACTTCGTTTCCG<br>TCTTGGCTACCACGCCATTCCATCTATGAGTCATGTACA<br>CCTGCATGTTATTTCGCAAGACTTTGATTCGCCATGCTT<br>GAAAAACAAGAAGCACTGGAATAGTTTCAACACAGAATA<br>TTTTCTTGAGTCCCAAGCCGTTATCGAGATGGTACAGGA<br>GGCTGGACGTGTTACTGTGCGTGACGGGATGCCCGAA<br>CTTCTGAAACTGCCACTTCGTTGTCACGAGTGTCAGCAG<br>TTATTACCCTCCATCCCTCAGTTGAAAGAACATCTGCGC<br>AAACACTGGACTCAGTGTACCCAGCTTT | In cells translation (Figure 4I) |
